# Iron-sulfur cluster repair contributes to *Y. pseudotuberculosis* survival within deep tissues

**DOI:** 10.1101/576454

**Authors:** Kimberly M. Davis, Joanna Krupp, Stacie Clark, Ralph R. Isberg

## Abstract

To successfully colonize host tissues, bacteria must respond to and detoxify many different host-derived antimicrobial compounds, such as nitric oxide (NO). NO has direct antimicrobial activity through attack on iron-sulfur (Fe-S) cluster-containing proteins. NO detoxification plays an important role in promoting bacterial survival, but it remains unclear if repair of Fe-S clusters is also important for bacterial survival within host tissues. Here we show that the Fe-S cluster repair protein, YtfE, contributes to the survival of *Y. pseudotuberculosis* within the spleen following nitrosative stress. *Y. pseudotuberculosis* forms clustered centers of replicating bacteria within deep tissues, where peripheral bacteria express the NO-detoxifying gene, *hmp*. *ytfE* expression also occurred specifically within peripheral cells at the edges of microcolonies. In the absence of *ytfE*, the area of microcolonies was significantly smaller than WT, consistent with *ytfE* contributing to the survival of peripheral cells. The loss of *ytfE* did not alter the ability of cells to detoxify NO, which occurred within peripheral cells in both WT and Δ*ytfE* microcolonies. In the absence of NO-detoxifying activity by *hmp*, NO diffused across Δ*ytfE* microcolonies, and there was a significant decrease in the area of microcolonies lacking *ytfE*, indicating that *ytfE* also contributes to bacterial survival in the absence of NO detoxification. These results indicate a role for Fe-S cluster repair in the survival of *Y. pseudotuberculosis* within the spleen, and suggest that extracellular bacteria may rely on this pathway for survival within host tissues.

## Introduction

Nitric oxide (NO) is a diffusible gas that has a wide range of physiological functions within mammals (1, 2). The effects of NO are tissue concentration-dependent, as it promotes vasodilation, cell proliferation and cell differentiation at low concentrations (3, 4), while high concentrations drive apoptosis and defense against bacterial, fungi, and parasites (5-8). NO is produced by three different nitric oxide synthase (NOS) isoforms within mammalian tissues: the neuronal NOS (nNOS), endothelial NOS (eNOS), and inducible NOS (iNOS). nNOS and eNOS are expressed at low levels by endothelial and neuronal cells respectively (2). In contrast, iNOS is expressed by a wide range of cell types, specifically in response to NF-κB-dependent sensing of, and is responsible for the high levels of NO produced during infection (1, 9).

NO has direct antimicrobial activity and can also react with reactive oxygen species to produce additional toxic compounds, such as peroxynitrite. NO may be bacteriostatic, while peroxynitrite is known to have direct bactericidal activity (10). NO antibacterial activity occurs through nitrosylation of iron-sulfur (Fe-S) cluster-containing proteins, which play critical roles in cellular respiration, DNA synthesis, and gene regulation. NO also targets heme groups, reactive thiols, tyrosyl radicals, and can cause DNA damage (1, 10). One of the global regulators in *E. coli* that has been shown to be inactivated by NO is NsrR (11), which regulates the response to nitrosative stress and is also associated with the oxidative stress response (12-14). Nitrosylation of the NsrR-associated Fe-S cluster relieves repression of at least 60 genes in *E. coli* (15-18). Included in this regulon is the *hmp* gene, which encodes a flavohemoglobin that detoxifies NO (19, 20), and YtfE (also known as Repair of Iron Centers (RIC) in *E. coli*), which functions to repair Fe-S clusters following NO damage (13, 17). Repair of Fe-S cluster proteins by YtfE can eliminate the need for new Fe-S cluster biogenesis when Fe availability is limited (13, 21, 22). A variety of studies argue that YtfE contributes to bacterial survival within host cells, but it remains unclear if YtfE contributes to the survival of extracellular pathogens replicating within host tissues, and if YtfE contributes to survival following exposure to NO (23-25).

*Yersinia pseudotuberculosis* is an oral pathogen that is typically contained within intestinal tissues and gut-associated lymphoid tissues, but has the capacity to spread systemically in susceptible individuals (26-29). Following bloodstream access, *Y. pseudotuberculosis* colonizes deep tissue sites where individual bacteria replicate to form clonal microcolonies (30-32). Neutrophils, monocytes, and macrophages are recruited to sites of bacterial replication, which are kept at bay by *Yersinia*, primarily via substrates of the bacterial type III secretion system (33-35). Recruited monocytes and macrophages produce NO, which diffuses across a layer of neutrophils and is inactivated at the periphery of the microcolony by bacteria expressing Hmp, preventing diffusion of NO into the interior of the microcolony (32). The consequences of the selective attack of NO on the peripheral bacteria are unclear, although it is likely that nitrosative stress may slow their growth. Additional members of the nitrosative stress response, such as YtfE, may also be required in peripheral cells to ensure their survival.

Bacteria responding to RNS appear to recover and remain viable, consistent with members of the NsrR regulon cooperating to repair NO-mediated damage in peripheral bacteria within microcolonies. Two of the members of the *E. coli* NsrR regulon, *hmp* and *ytfE,* are upregulated in the bubonic plague model of *Yersinia pestis* infection and during Peyer’s patch colonization by *Y. pseudotuberculosis* (36, 37). The similar expression patterns of *hmp* and *ytfE* may suggest that both genes are also members of the NsrR regulon in *Yersinia,* and could both contribute to microbial fitness during microcolony growth. Additionally, very few studies have explored the role of Fe-S cluster repair during infection with extracellular bacteria. Here, we show that *ytfE* contributes to the survival of extracellular bacteria, specifically through upregulation of *ytfE* within peripheral cells of *Y. pseudotuberculosis* microcolonies.

## Results

### *Y. pseudotuberculosis ytfE* expression is regulated by NsrR and occurs within peripheral cells during growth in the spleen

The *ytfE* gene is known to be a member of the NsrR regulon in a number of bacterial species (13, 14, 38), so *ytfE* is expected to be repressed by NsrR in *Y. pseudotuberculosis*. We also expected that *ytfE* is transcribed during *Y. pseudotuberculosis* growth within the mouse spleen as the NsrR-regulated *hmp* gene is expressed in this tissue (23-25, 32, 36). To determine if *Y. pseudotuberculosis ytfE* is expressed during splenic growth, C57BL/6 mice were intravenously challenged with bacteria and bacterial RNA was isolated at day 3 post-inoculation (PI). Based on qRT-PCR analysis, *ytfE* transcript levels increased within the mouse spleen relative to the inoculum culture grown in the absence of an NO-generating system, with marked mouse-to-mouse variation (Figure 1A) (32). To determine if *ytfE* expression is NsrR-dependent, we compared *ytfE* transcription levels in WT and Δ*nsrR* strains in the presence (+) and absence (-) of nitrogen stress imparted by acidified nitrite (NO_2_). Transcription of *ytfE* increased by over 100X in the WT strain with the addition of nitrogen stress (Figure 1B). Transcription of *ytfE* was high in the Δ*nsrR* strain in either the presence or absence of nitrogen stress, indicating that NsrR negatively regulates *ytfE* transcription. A slight increase in *ytfE* expression in the Δ*nsrR* strain with nitrogen stress suggests additional pathways may regulate *ytfE* expression.

**Figure 1:**
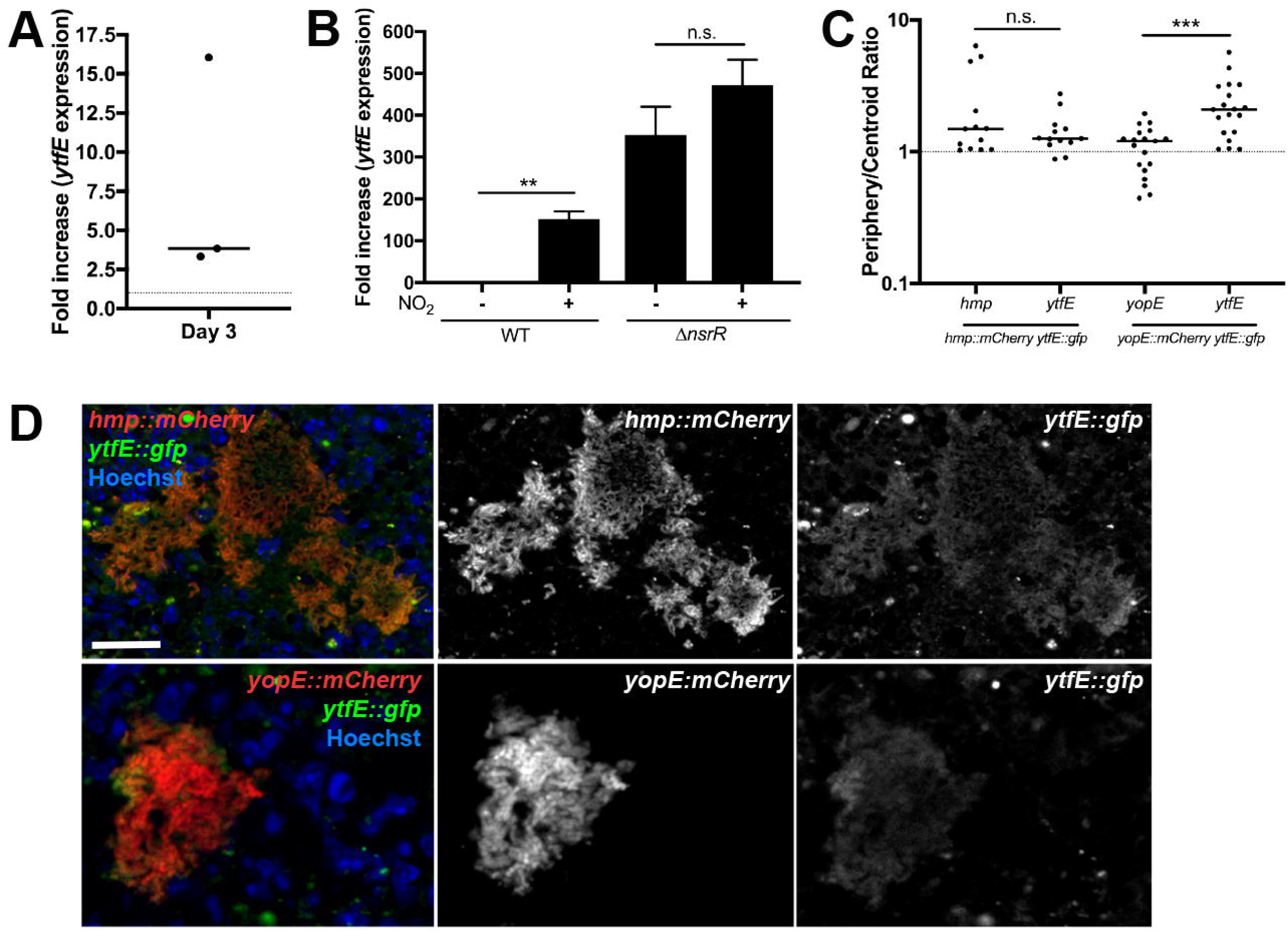
*ytfE* expression is regulated by NsrR and occurs within peripheral cells. A) C57BL/6 mice were inoculated intravenously (i.v.) with 10^3^ WT *Y. pseudotuberculosis*, and spleens were harvested at day 3 post-inoculation (PI). Bacterial transcripts were isolated from splenic tissue, transcript levels were quantified by qRT-PCR relative to 16S, and the fold increase in *ytfE* transcript levels is shown relative to the inoculum. Each dot represents an individual mouse. B) Nitrogen stress was induced (NO_2_, +) in cultures of WT and Δ*nsrR* strains, and compared to untreated cultures (NO_2_, -). The *ytfE* transcript levels are expressed relative to 16S, fold increase is relative to the average level in untreated WT cultures. Each column: 5 biological replicates, mean and SEM. C) C57BL/6 mice were inoculated i.v. with the WT *hmp::mCherry ytfE::gfp* strain or the WT *yopE::mCherry ytfE;:gfp* strain, and spleens were harvested at day 3 PI for fluorescence microscopy. Fluorescent signal intensity was determined within the same peripheral and centroid cells for *hmp* and *ytfE* reporters or *yopE* and *ytfE* reporters, and divided to generate the periphery/centroid ratio (4 mice/group). Dots: individual microcolonies. D) Representative image of *hmp::mCherry* and *ytfE::gfp* reporters (top panels) or *yopE::mCherry* and *ytfE::gfp* reporters (bottom panels); merge and single channel images are shown. Scale bars: 20µm. Statistics: Wilcoxon matched pairs, **p<0.01, n.s.: not significant.

Hmp, another member of the NsrR regulon, is specifically expressed on the periphery of *Y. pseudotuberculosis* microcolonies during splenic growth (32). At the transcriptional level, this is associated with considerable variation in expression levels between individual mice. Variation is likely due to the presence of different size microcolonies within different organs, as distinct peripheral expression of *hmp* is visualized in large microcolonies, and smaller microcolonies are more homogenous. The variable *ytfE* expression levels in the mouse and NsrR-dependence is reminiscent of *hmp* expression, indicating that there could be a link between spatial expression and inter-mouse variation. To determine if *ytfE* and *hmp* have similar expression patterns during growth in the spleen, mice were intravenously inoculated with a WT *Y. pseudotuberculosis* strain containing *hmp::mCherry* (chromosomal integration of *mCherry* downstream of *hmp*) and *ytfE::gfp* (chromosomal integration of *gfp* downstream of *ytfE*). Microcolonies were visualized within the spleen using fluorescence microscopy, and *ytfE* and *hmp* reporter signals were quantified within the same cells at the center and periphery of the microcolonies (Materials and Methods). The *hmp* reporter signal increased within individual cells at the periphery relative to cells at the centroid of microcolonies, which generated a ratio value greater than 1, consistent with *hmp* peripheral expression (Figure 1C). The *ytfE* signal was dim relative to *hmp*, but also increased at the periphery of microcolonies relative to the centroid. The Periphery/Centroid signal intensity ratio value for *ytfE* was similar to *hmp*, indicating that *ytfE* was expressed at the periphery of microcolonies (Figure 1C). To confirm the dim *ytfE* signal was not due to mCherry fluorescence detected in the gfp channel, experiments were also performed in a WT strain containing *yopE::mCherry* and *ytfE::gfp*. *yopE* and *ytfE* exhibited distinct reporter patterns indicating the detected gfp signal was due to *ytfE* reporter expression (Figure 1C, 1D). There was significant overlap in the *hmp* and *ytfE* signals within these images, and based on NsrR-dependent regulation of both genes, it is likely that *hmp* and *ytfE* were expressed in the same cells (Figure 1D).

### *ytfE* contributes to the virulence of *Y. pseudotuberculosis* in the spleen

YtfE repairs Fe-S clusters damaged by NO, and there appears to be sufficient NO at the periphery of splenic microcolonies to allow synthesis of this protein and to promote repair within this subpopulation of bacteria. To determine if loss of *yftE* alters the overall fitness of *Y. pseudotuberculosis* during growth within the spleen, we constructed a *Y. pseudotuberculosis* strain that lacks the *ytfE* gene (Δ*ytfE*) and harbors a constitutive *gfp*-expressing plasmid, to visualize growth within the spleen. To compare differences in relative fitness, mice were infected intravenously with equal amounts of WT mCherry^+^ and Δ*ytfE* GFP^+^ strains, and spleens were harvested at day 3 PI, a late stage of infection in this model, to determine the competitive index by colony forming units (CFUs) and quantify microcolony areas within the same animals. At day 3 PI, the median competitive index was significantly less than 1, indicating lowered fitness of the Δ*ytfE* strain relative to the WT strain (Figure 2A; p=0.0115). The areas of individual microcolonies within these co-infected tissues were also visualized and quantified by fluorescence microscopy in 11 mice. The Δ*ytfE* microcolonies were significantly smaller than WT microcolonies within the same organs, indicating that *ytfE* contributes to the survival of *Y. pseudotuberculosis* in the spleen, presumably due to lowered fitness of the bacterial population located at the periphery of the microcolonies (Figure 2B).

**Figure 2:**
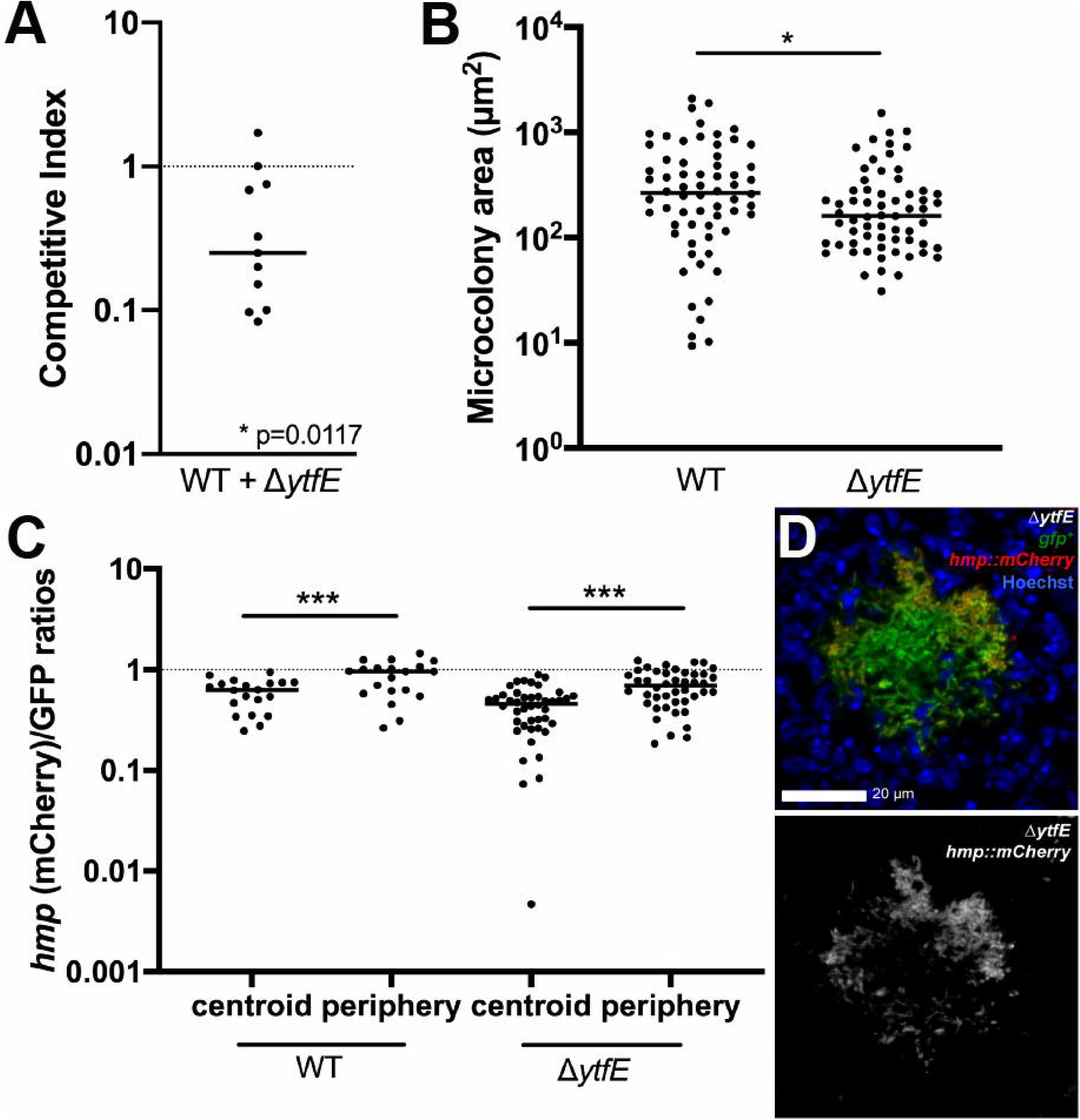
*ytfE* contributes to growth in the spleen. A) Co-infection with the WT and *ΔytfE* strains. C57BL/6 mice were inoculated i.v. with an equal mixture of mCherry^+^ (*yopE::mCherry*) WT and GFP^+^ Δ*ytfE* strains, and spleens were harvested at day 3 PI. Competitive index: ratio of Δ*ytfE*/WT CFUs in spleen at day 3, divided by ratio of Δ*ytfE*/WT CFUs in the inoculum. Dots: individual mice. Dotted line: equal fitness, replicates completed on two separate days. B) WT and Δ*ytfE* microcolony areas (µm^2^) from the co-infection in panel 2A quantified (Experimental Procedures) in 11 mice. C) C57BL/6 mice were inoculated i.v. with the WT GFP^+^ *hmp::mCherry* or *ΔytfE* GFP^+^ *hmp::mCherry* strains, and spleens were harvested at day 3 PI for fluorescence microscopy. Reporter signals were quantified within peripheral and centroid cells, *hmp* reporter signal was divided by GFP. D) Representative image of a *ΔytfE* GFP^+^ *hmp::mCherry* microcolony, stained with Hoechst to detect host nuclei; merge and *hmp* single channel images are shown Statistics: A): Wilcoxon Signed Rank Test, compared to a value of 1, B): Mann-Whitney, C): Wilcoxon matched pairs, *p<0.05, ***p<0.001.

Since YtfE could directly repair the Fe-S cluster of NsrR, the absence of *ytfE* could alter expression of the NsrR regulon, by resulting in heightened expression within peripheral cells. To determine if microcolonies from the *Y. pseudotuberculosis* Δ*ytfE* strain have sustained expression of the NsrR regulon relative to the WT strain, we infected mice intravenously with WT GFP^+^ *hmp::mCherry* or Δ*ytfE* GFP^+^ *hmp::mCherry* integrated *hmp* reporter strains, and spleens were harvested at day 3 PI to visualize reporter expression by fluorescence microscopy. Similar to Fig. 1D, *hmp* reporter expression was significantly higher at the periphery relative to the centroid in the WT strain (Figure 2C). The reporter expression pattern was very similar in Δ*ytfE* microcolonies, indicating that there was still a gradient of NO exposure in the mutant, in which Hmp activity in the peripheral population protects the central population of bacteria, and that loss of *ytfE* did not significantly alter expression of the NsrR regulon (Figure 2D).

### *ytfE* contributes to bacterial survival in the absence of *hmp*

YftE contributed to the growth of *Y. pseudotuberculosis* microcolonies in the spleen despite expression limited to the microcolony periphery. We were then interested in determining if YtfE-mediated repair played an important role in the context of a Δ*hmp* strain, where all bacteria in a microcolony are exposed to NO (32). To address this point, we challenged mice intravenously with Δ*hmp* GFP^+^ *P_hmp_::mCherry* and Δ*hmp* Δ*ytfE* GFP^+^ *P_hmp_::mCherry* strains, and spleens were harvested at day 3 PI to quantify CFUs, visualize microcolony areas, and detect reporter signals by fluorescence microscopy. The Δ*hmp* Δ*ytfE* strain showed no potentiation of the defect in single strain infections, but this defect can be observed during co-infection. The median competitive index (CI) value for the double mutant was below 1, but was not significantly less than 1, which suggests that Δ*hmp* Δ*ytfE* strain may not be less fit than Δ*hmp* (Figure 3A). Interestingly, in 4 mice the CI was at least 1, indicating that the Δ*hmp* Δ*ytfE* strain could compete efficiently with Δ*hmp* in these animals. We then compared the total CFU in the organs in mice in which the Δ*hmp* strain outcompeted Δ*hmp* Δ*ytfE*, or in mice in which no such outcompetition took place (Figure 3B). The total CFUs were significantly lower in organs in which the Δ*hmp* did not outcompete the Δ*hmp*Δ*ytfE* strain, indicating that the fitness differences were suppressed in animals showing increased restriction of bacterial growth. It is also possible that the Δ*hmp*Δ*ytfE* strain had reduced levels of initial seeding within tissues. The microcolony areas were quantified within all the spleens depicted in Figure 3A, and the areas of Δ*hmp* Δ*ytfE* microcolonies were significantly smaller than Δ*hmp* (Figure 3C). Δ*hmp* Δ*ytfE* microcolonies had similar *P_hmp_* reporter expression at the centroid and periphery, indicating NO diffused across these centers (Figure 3D). Together, these results confirm that *ytfE* contributes to bacterial survival in the absence of Hmp detoxifying activity.

**Figure 3:**
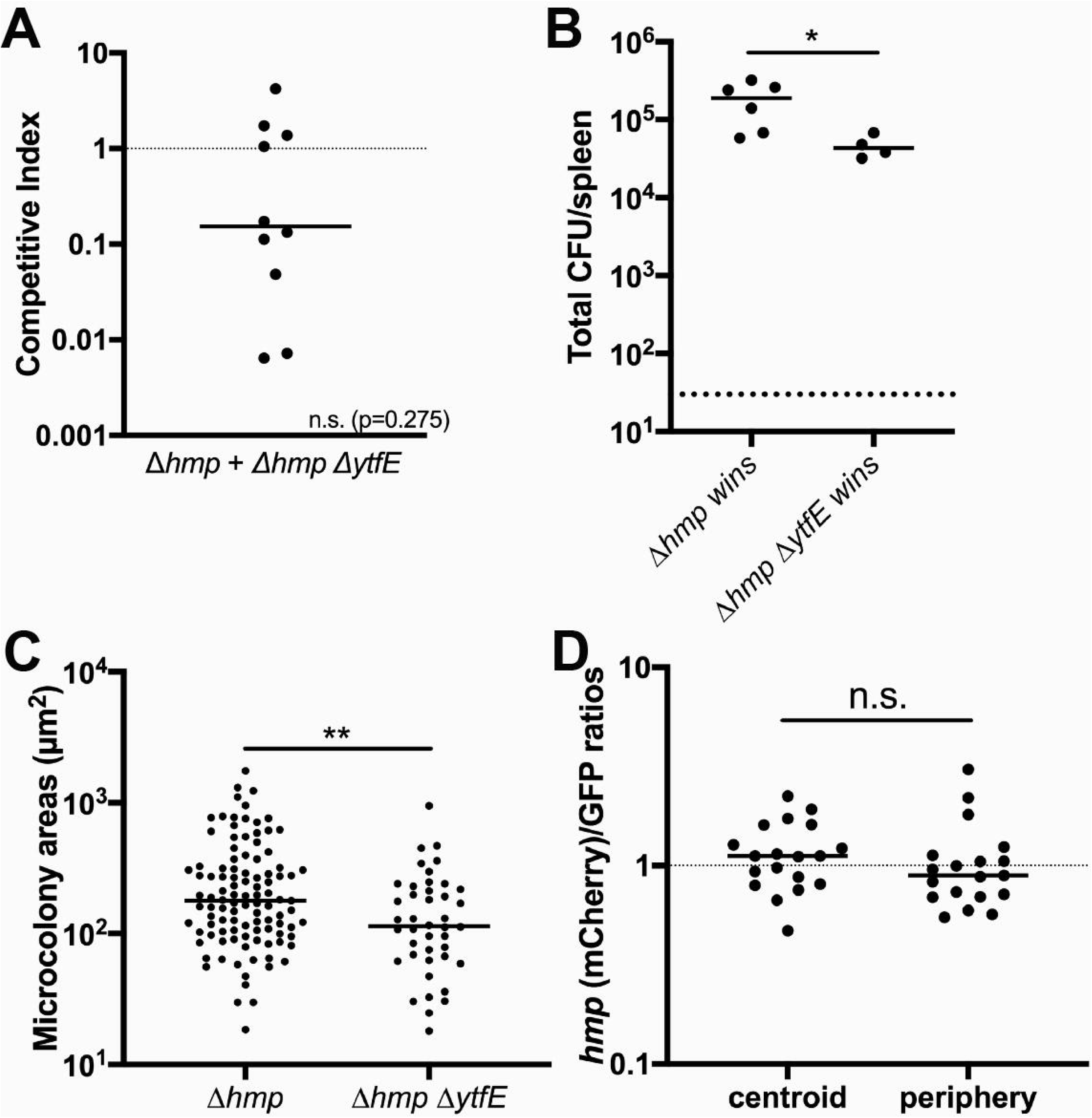
Δ*ytfE* contributes to survival in the absence of *hmp*. A) C57BL/6 mice were inoculated i.v. with an equal mixture of *hmp* GFP^+^ and Δ*hmp* Δ*ytfE* GFP^+^ *P_hmp_::mCherry* strains, and spleens were harvested at day 3 PI. Competitive index: ratio of Δ*hmp* Δ*ytfE*/Δ*hmp* CFUs in spleen at day 3, divided by ratio of Δ*hmp* Δ*ytfE*/Δ*hmp* CFUs in the inoculum. Dots: individual mice. Dotted line: represents equal fitness. Statistics: Wilcoxon Signed Rank Test, compared to a value of 1, n.s.: not significant. B) Total CFUs in the spleen during co-infection, when CI was less than 1 (Δ*hmp* wins) or above or equal to 1 (Δ*hmp* Δ*ytfE* wins). Dots: individual mice. C) Δ*hmp* and Δ*hmp* Δ*ytfE* microcolony areas (µm^2^) from the co-infection quantified (Experimental Procedures) in 10 mice. Dots: individual microcolonies. D) Reporter signals were quantified within peripheral and centroid cells in the Δ*hmp* Δ*ytfE* strain during co-infection, *hmp* reporter signal was divided by GFP. Dots: individual microcolonies. Statistics: B) and C): Mann-Whitney, D): Wilcoxon matched pairs, *p<0.05, **p<0.01, n.s.: not significant.

### *ytfE* has limited effects on NO sensitivity in the absence of *hmp*

The Δ*ytfE* single mutant strain had reduced fitness relative to the WT strain (Figure 2), however the single Δ*ytfE* mutant was not significantly more sensitive than the WT strain to the presence of acidified nitrite during growth in culture. Similarly, despite the lowered fitness of the Δ*hmp* Δ*ytfE* strain in spleens relative to Δ*hmp*, the Δ*hmp* Δ*ytfE* strain was not significantly more sensitive than the Δ*hmp* mutant strain to the presence of acidified nitrite during growth in culture (Figure 4A; compare +NO_2_ samples). The absence of *ytfE* also did not significantly alter the sensitivity of strains to the NO donor compound, DETANONOate, during growth in minimal media (Figure 4B). This is consistent with previous reports in other organisms (38, 39), perhaps due to the presence of other backup repair pathways that are active in the absence of Hmp or because YtfE plays a role in protection from other stress species.

**Figure 4:**
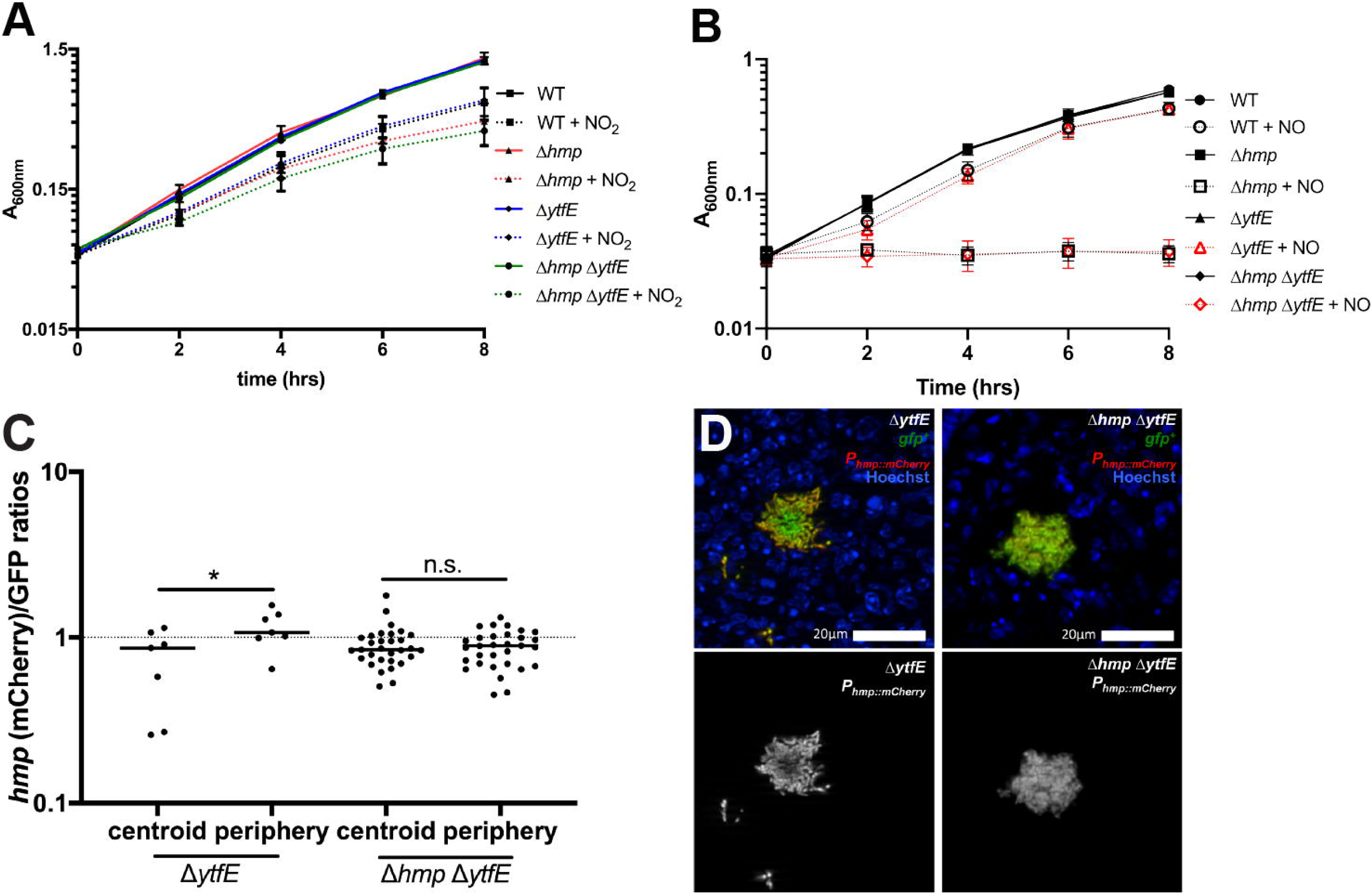
*ytfE* has limited effects on NO sensitivity in the absence of *hmp.* A) Optical density (A_600nm_) was measured every 2 hours (time, hrs) within cultures of the indicated strains during incubation in LB pH5.5 in the presence (+NO_2_) and absence of NaNO_2_. Data represents two independent experiments. B) Optical density (A_600nm_) was measured every 2 hours (time, hrs) within cultures of the indicated strains during incubation in M9 minimal media in the presence (+NO) and absence of the DETANONOate NO donor. Data represents three independent experiments. C) C57BL/6 mice were inoculated i.v. with the Δ*ytfE* GFP^+^ *P_hmp_::mCherry* or Δ*hmp ΔytfE* GFP^+^ *P_hmp_::mCherry* strains, and spleens were harvested at day 3 PI for fluorescence microscopy. Reporter signals were quantified within peripheral and centroid cells, *hmp* reporter signal was divided by GFP. Dots: individual microcolonies. D) Representative images of Δ*ytfE* and *Δhmp ΔytfE* microcolonies; merge and *hmp* single channel images are shown. Statistics: Wilcoxon matched pairs, *p<0.05, n.s.: not significant.

We then compared *hmp* reporter expression in Δ*ytfE* and Δ*hmp*Δ*ytfE* strains to confirm that NO diffusion occurred across Δ*hmp* Δ*ytfE* microcolonies using plasmid-borne reporters. Mice were infected intravenously with Δ*ytfE* GFP^+^ *P_hmp_::mCherry* or Δ*hmp* Δ*ytfE* GFP^+^ *P_hmp_::mCherry* strains, and spleens were harvested 3 days PI to visualize reporter expression by fluorescence microscopy. The Δ*ytfE* strain had peripheral *P_hmp_* reporter expression, as seen with the chromosomally-integrated *hmp* reporter, indicating that NO diffusion across the microcolony is inhibited by peripheral cells in Δ*ytfE* microcolonies (Figure 4C, 4D). Δ*hmp* Δ*ytfE* microcolonies showed no such preference for the periphery, indicating NO diffused across these centers, as expected based on the loss of *hmp*.

### Rescue of *ytfE* restores microcolony size

To show that the loss of *ytfE* was responsible for decreased microcolony size, we rescued the Δ*ytfE* strain with a WT copy of *ytfE*, and transformed the *ytfE* rescued strain with the constitutive *gfp*-expressing plasmid. Mice were infected intravenously with WT mCherry^+^ and *ytfE* rescued GFP^+^ strains, and spleens were harvested at day 3 PI to quantify CFUs and visualize microcolony areas by fluorescence microscopy. The median competitive index for the *ytfE* rescued strain was close to a value of 1, indicating the fitness of this strain was roughly equivalent to the WT strain (Figure 5A). The microcolony areas were quantified within all the spleens depicted in Figure 5A, and the areas of *ytfE* rescued microcolonies were very similar to WT microcolonies (Figure 5B, 5C). These results indicate that the Δ*ytfE* strain was rescued by the WT copy of *ytfE*, which confirms that the reduced fitness of the Δ*ytfE* strain was specifically due to a loss of *ytfE*.

**Figure 5:**
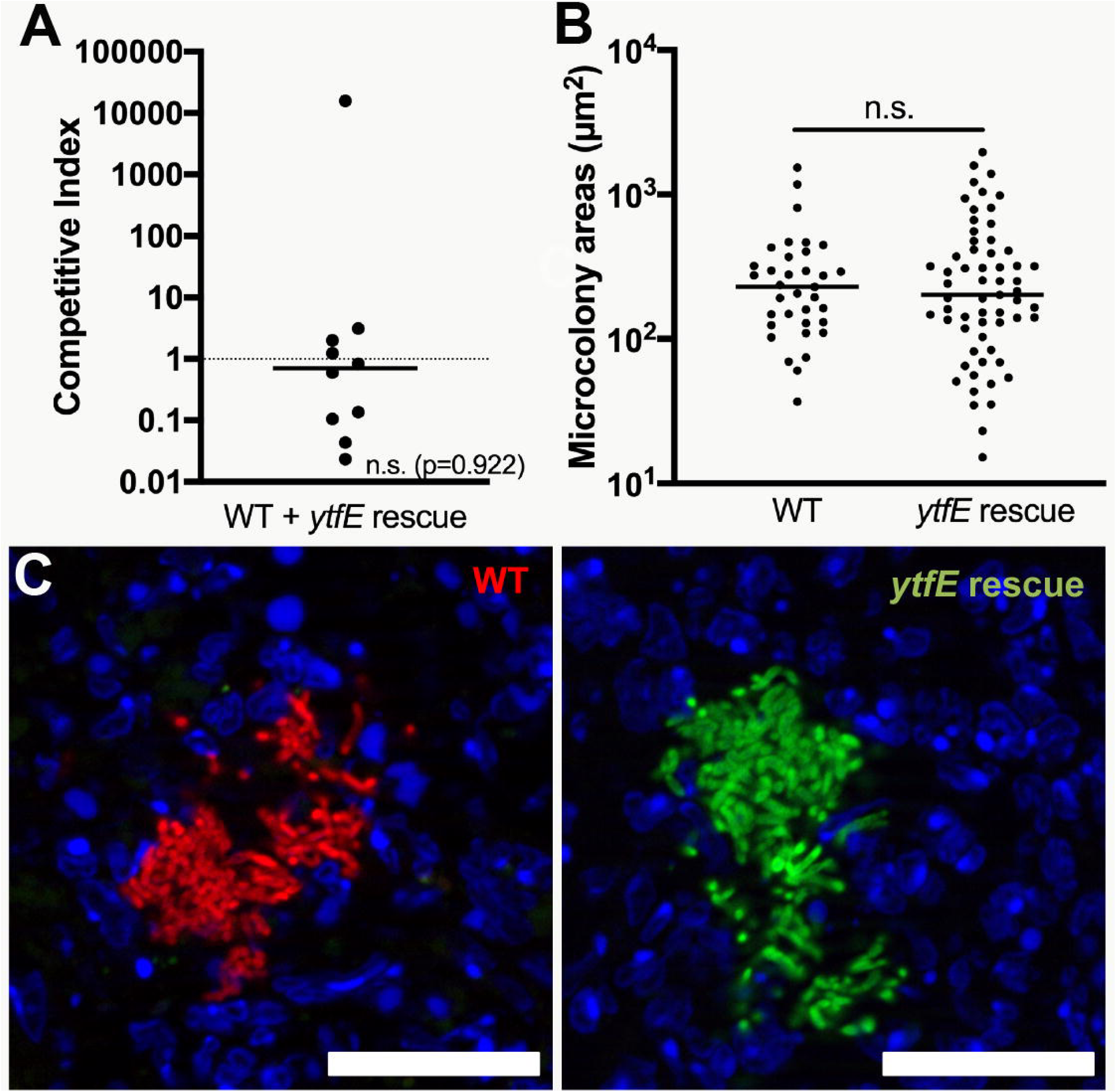
Rescue of *ytfE* restores fitness of the Δ*ytfE* strain. A) C57BL/6 mice were inoculated i.v. with an equal mixture of *ytfE* rescued GFP^+^ and WT mCherry^+^ strains, and spleens were harvested at day 3 PI. Competitive index: ratio of *ytfE* rescue/WT CFUs in spleen at day 3, divided by ratio of *ytfE* rescue/WT CFUs in the inoculum. Dots: individual mice. Dotted line: equal fitness, replicates completed on two separate days. Statistics: Wilcoxon Signed Rank Test, compared to a value of 1, n.s.: not significant. B) WT and *ytfE* rescue microcolony areas (µm^2^) from the co-infection quantified (Experimental Procedures) in 10 mice. Dots: individual microcolonies. C) Representative images of WT mCherry^+^ and *ytfE* rescued GFP^+^ microcolonies. Scale bar: 20µm. Statistics: Mann-Whitney, n.s.: not significant.

## Discussion

The detoxification of NO and other reactive nitrogen species is critical for bacterial survival within host tissues (40-43). Bacterial proteins involved in NO detoxification, however, are not synthesized until NO accumulates and damages the Fe-S clusters of NsrR, resulting in inactivation of this repressor (15, 16). Many additional Fe-S cluster-containing proteins are simultaneously damaged, so bacteria need to either repair damaged proteins or synthesize replacement proteins, while simultaneously replenishing proteins that detoxify NO and prevent further damage (17, 38). Although Fe-S cluster repair is likely required for bacterial survival, it has been unclear whether or not this plays an important role within host tissues. We have chosen to ask this question in a mouse model of *Y. pseudotuberculosis* infection, where it is known that bacteria respond to RNS and that Fe concentrations are limiting (24, 25, 32). *ytfE* expression had been detected in *Yersinia* species replicating within host tissues, however it was unclear if Fe-S cluster repair or assembly play a significant role during infection (36, 37). Here, we have shown that Fe-S cluster repair contributes to successful *Y. pseudotuberculosis* replication within the spleen.

We found that a place where Fe-S cluster repair in *Y. pseudotuberculosis* likely occurs is in the peripheral subpopulation of bacteria that responds to NO assault within microcolonies. This is consistent with a role for YftE in supporting survival of the peripheral subpopulation. In the absence of *ytfE*, we would expect bacteria on the periphery to be exposed to stress associated with NO exposure, leading to sequential loss of the peripheral population and progressively smaller microcolonies. Consistent with this hypothesis, we observed Δ*ytfE* microcolonies were smaller than those established by the WT strain. It is expected that the difference between WT and Δ*ytfE* microcolony areas will become progressively more pronounced as the infection proceeds, because Δ*ytfE* microcolonies should continuously lose their peripheral subpopulation. This prediction is based on our previous observation that NO-sensitive microcolonies are progressively reduced during the course of disease, with elimination of Δ*hmp* bacteria by NO most pronounced at late timepoints post-inoculation, concurrent with a timepoint in which the animals inoculated with the WT strain are moribund (32).

NO alone was not sufficient to limit the growth of Δ*ytfE* bacteria in bacteriological media, consistent with previous studies that indicated the uropathogenic *E. coli* (UPEC) Δ*ytfE* strain had reduced intracellular survival within host cells, but was not sensitive to exogenous NO alone (39). Presumably, the Δ*ytfE* strain is sensitive to other antimicrobial compounds generated within host tissues, possibly by the generation of a variety of reactive nitrogen species (RNS) as a consequence of NO reaction with reactive oxygen species (ROS) or other compounds. Upregulation of *ytfE* following hydrogen peroxide-mediated damage of *Staphylococcus aureus* indicates that YtfE may play a broad role in repair, instead of just a response to nitrogen stress (44, 45). Additional studies in *E. coli* also suggest that YtfE repairs Fe-S cluster proteins damaged by hydrogen peroxide (13, 46-49).

An additional regulatory signal for the *yftE* gene is iron limitation, which depresses synthesis of new Fe-S cluster proteins, thus requiring cellular YftE function (14, 21, 50). The spleen is expected to be an iron-limiting environment requiring Fe-S cluster repair, but we found no evidence for NO-independent induction of the *ytfE* gene within the center of microcolonies, using our fluorescent promoter constructions. This contrasts with data arguing that *Y. pseudotuberculosis* expresses many Fe acquisition genes during growth in host tissues, consistent with Fe-limiting conditions within host tissues (25, 37). It is possible that the center of the microcolony represents a protected environment with low exposure to stresses such as reactive nitrogen and oxygen species as well as limited damage to Fe-S centers. Additionally, *ytfE* expression may be induced at very low NO concentrations, as seen in *Salmonella enterica* serovar Typhimurium (38), but may require severe iron limitation for expression in the absence of NO.

The loss of YftE function also has the potential to alter metabolite levels due to disruption of protein functions that are Fe-S center-related. In the presence of RNS, a Δ*ytfE* strain may have reduced aconitase and fumarase activity due to the role of YtfE in Fe-S cluster repair specifically for these proteins (47). The Fe-S cluster of the NsrR repressor is also repaired by YtfE, which could lead to prolonged expression of the NsrR regulon in the presence of NO (13). Our results indicate that expression of the NsrR regulon is similar in the presence or absence of *ytfE*, although it remains possible that prolonged expression of the NsrR regulon could be detected at later timepoints post-inoculation. Future work will investigate these issues, and determine the interplay between iron regulation, NO-induced damage and repair of critical Fe-S centers.

## Materials and Methods

### Bacterial strains & growth conditions

The WT *Y. pseudotuberculosis* strain, YPIII, was used throughout. For all mouse infection experiments, bacteria were grown overnight to post-exponential phase in 2xYT broth (LB, with 2x yeast extract and tryptone) at 26o C with rotation. Exponential phase cultures were sub-cultured 1:100 from overnight cultures, and grown at 26o C with rotation for an additional 2 hours. Sodium nitrite (2.5mM) was added to LB pH5.5 to induce the nitrogen stress response. DETA-NONOate (2.5mM) NO donor compound (Cayman Chemicals) was added to M9 minimal media to test NO sensitivity.

### Generation of ytfE mutant strains

The *hmp* and *nsrR* deletion strains were previously described (32). Deletion derivative strains were generated for *ytfE* by amplifying the start codon + 3 downstream codons, the 3’ terminal 3 codons + the stop codon and fusing these fragments to generate a start + 6 aa + stop construct. Deletion constructions were amplified with 500 base pairs flanking sequence on each side, cloned into the suicide vector, pSR47S, and transformed into *Y. pseudotuberculosis*. Sucrose selection was used to select for bacteria that had incorporated the desired mutation after a second cycle of recombination (31). PCR, sequencing, and qRT-PCR were used to confirm deletion strains.

### Integrated ytfE reporter construction (ytfE^+^rescue strain)

The *ytfE::gfp* reporter was generated by cloning *gfp* immediately downstream of the *ytfE* gene (in between the *ytfE* stop codon and terminator sequence) by overlap extension PCR. The *ytfE* start codon was amplified with 500 base pairs upstream flanking sequence, while the stop codon of *gfp* was amplified with 500 base pairs downstream flanking sequence. This fragment was cloned into the suicide vector, pSR47S, and transferred by conjugation into WT *Y. pseudotuberculosis hmp::mCherry* (chromosomally integrated), selecting for kanamycin resistance. For the *ytfE^+^* rescue strain, a WT *ytfE* gene product, including 500 base pairs upstream and downstream of *ytfE*, was amplified from genomic DNA and cloned into pSR47S. This vector was transferred by conjugation into Δ*ytfE Y. pseudotuberculosis*, selecting for kanamycin resistance. A second round of recombination was selected on sucrose-containing media to isolate strains that had recombined each *ytfE* construct. PCR and sequencing were used to confirm integration of *gfp* or rescue of the *ytfE* deletion.

### Generation of plasmid-based reporter strains

Several of the *Y. pseudotuberculosis* reporter strains in this study have been previously described: WT GFP^+^, WT mCherry^+^ (*yopE::mCherry)*, WT *hmp::mCherry*, WT GFP^+^ *hmp::mCherry*, and Δ*hmp* GFP^+^ (32). For this study, GFP^+^ strains were constructed by transforming deletion strains with the constitutive GFP plasmid, which expresses GFP from an unrepressed *P_tet_* of pACYC184. The *P_hmp_::mCherry* was also transformed into GFP^+^ strains. The *P_hmp_::mCherry* transcriptional fusion was previously constructed by fusing the *hmp* promoter to *mCherry* using overlap extension PCR, and cloned into the pMMB67EH plasmid (32).

### Murine model of systemic infection

Six to 8-week old female C57BL/6 mice were obtained from Jackson Laboratories (Bar Harbor, ME). All animal studies were approved by the Institutional Animal Care and Use Committee of Tufts University. Mice were injected intravenously with 10^3^ bacteria for all experiments. For co-infection experiments, mice were inoculated with 5 x 10^2^ CFU of each strain, for a total of 10^3^ CFUs. At the indicated timepoints post-inoculation (PI) (3 days), spleens were removed and processed.

### qRT-PCR to detect bacterial transcripts in broth-grown cultures

Bacterial cells were grown in broth to A_600_ = 0.3 under appropriate stress conditions, pelleted, resuspended in Buffer RLT (QIAGEN) + ß-mercaptoethanol, and RNA was isolated using the RNeasy kit (QIAGEN) according to manufacturer’s protocol. DNA contamination was eliminated using the DNA-free kit (Ambion) according to manufacturer’s protocol. RNA was reverse transcribed using M-MLV reverse transcriptase (Invitrogen), in the presence of the RNase inhibitor, RnaseOUT (Invitrogen), according to manufacturer’s protocol. Approximately 30 ng cDNA was used as a template in reactions with 0.5 µM of forward and reverse primers and SYBR Green (Applied Biosystems) according to manufacturer’s protocol. Control samples were prepared that lacked M-MLV, to confirm DNA was eliminated from samples and was not amplified by qRT-PCR. Reactions were carried out using the StepOnePlus Real-Time PCR system, and relative comparisons were obtained using the ΔΔC_T_ or 2^-ΔCt^ method (Applied Biosystems).

### qRT-PCR to detect bacterial transcripts from mouse tissues

Mice were inoculated intravenously with the WT strain, and at day 3 PI spleens were harvested, and immediately submerged in RNALater solution (QIAGEN). Tissue was homogenized in Buffer RLT + ß-mercaptoethanol, and RNA was isolated using the RNeasy kit (QIAGEN) according to manufacturer’s protocol. Bacterial RNA was enriched following depletion of host mRNA and rRNA from total RNA samples, using the MICROB*Enrich* kit (Ambion) according to manufacturer’s protocol. DNA digestion, reverse transcription, and qRT-PCR were performed as described above.

### Fluorescence microscopy

C57BL/6 mice were inoculated intravenously with the *Y. pseudotuberculosis* WT strain, at day 3 PI spleens were harvested and immediately fixed in 4% paraformaldehyde in PBS for 3 hours. Tissues were frozen-embedded in Sub Xero freezing media (Mercedes Medical) and cut by cryostat microtome into 10µm sections. To visualize reporters, sections were thawed in PBS, stained with Hoechst at a 1:10,000 dilution, washed in PBS, and coverslips were mounted using ProLong Gold (Life Technologies). Tissue was imaged with 20x or 63x objectives, using a Zeiss Axio Observer.Z1 (Zeiss) fluorescent microscope with Colibri.2 LED light source, an Apotome.2 (Zeiss) for optical sectioning, and an ORCA-R^2^ digital CCD camera (Hamamatsu).

### Image analysis

Volocity image analysis software was used to quantify microcolony areas. Image J was used to quantify the signal intensity of each channel at the centroid and periphery of each microcolony, to generate relative signal intensities of fluorescent reporters. Thresholding was used to define the area of each microcolony, the centroid was calculated, and 0.01 pixel^2^ squares were selected to calculate values at the centroid. Peripheral measurements depict bacteria in contact with host cells.

## Author Contributions

Designed and performed experiments: KMD, JK, SC. Intellectual/conceptual contribution: KMD, RRI. Analyzed the data: KMD, JK, SC, RRI. Wrote the paper: KMD, RRI.

## Acknowledgments

We thank the members of the Isberg lab, who provided valuable advice and feedback throughout this project. We also thank the members of the Davis lab, who provided feedback and suggestions during the final steps of manuscript preparation. The authors of this manuscript declare no conflicts of interest. This work was supported by NIAID award R01 AI110684 as well as by an American Cancer Society-Ellison Foundation Postdoctoral Fellowship (PF-13-360-01-MPC), and a NIAID K22 Career Transition Award (1K22AI123465-01).

